# Foot strike pattern during running alters muscle-tendon dynamics of the gastrocnemius and the soleus

**DOI:** 10.1101/671370

**Authors:** Jennifer R. Yong, Christopher L. Dembia, Amy Silder, Rachel W. Jackson, Michael Fredericson, Scott L. Delp

**Affiliations:** Departments of Mechanical Engineering, Stanford University; Bioengineering, Stanford University; Orthopaedic Surgery, Stanford University

**Keywords:** rearfoot strike, forefoot strike, triceps surae, plantar flexors, Achilles tendon

## Abstract

Running is thought to be an efficient gait due, in part, to the behavior of the plantar flexor muscles and the elastic energy storage in the Achilles tendon. Although plantar flexor muscle mechanics and Achilles tendon energy storage have been explored during rearfoot striking, they have not been fully characterized during forefoot striking. This study examines how plantar flexor muscle-tendon mechanics during running differ between rearfoot and forefoot striking. We used musculoskeletal simulations, driven by joint angles and electromyography recorded from runners using both rearfoot and forefoot striking running patterns, to characterize plantar flexor muscle-tendon mechanics. The simulations revealed that foot strike pattern affected the soleus and gastrocnemius differently. For the soleus, forefoot striking resulted in decreased tendon energy storage and decreased positive fiber work compared to rearfoot striking. For the gastrocnemius, forefoot striking resulted in greater activation and increased negative fiber work compared to rearfoot striking. The increased activation and negative fiber work in the gastrocnemius during forefoot striking suggest that runners planning to convert to forefoot striking would benefit from a progressive eccentric gastrocnemius strengthening program to avoid injury.

## INTRODUCTION

The efficiency of running is enhanced by elastic energy storage in muscle-tendon units. This concept has supported the development of simple spring-mass models of human running^1^, which have been useful for understanding running mechanics^2^, predicting the energy cost of running^3^, and examining the effects of fatigue^4,5^. The plantar flexor muscles, in conjunction with the Achilles tendon, are major contributors to energy storage and return during running^6^, with energy being absorbed during early stance and released during late stance^7^. The elastic stretch and recoil of the Achilles tendon may contribute as much as 35% of the total energy storage and return during running^8^. Furthermore, the plantar flexor muscles are the largest contributors to body weight support and forward propulsion during running^9^.

The plantar flexor muscles and Achilles tendon span the ankle joint, suggesting that their mechanics would be affected by foot strike pattern. Rearfoot striking, characterized by landing on the heel, and forefoot striking, characterized by landing on the ball of the foot, are both naturally adopted foot strike patterns. Habitual rearfoot striking runners who transition to a forefoot striking pattern initially experience calf soreness^10,11^, indicating that altering foot strike pattern may affect the behavior of the plantar flexors and, therefore, energy storage in the Achilles tendon. Previous research has focused on understanding plantar flexor mechanics during rearfoot striking^12–15^, but how plantar flexor mechanics are affected by converting to forefoot striking is unclear.

Differences between rearfoot and forefoot striking suggest how plantar flexor muscle-tendon mechanics might be affected by foot strike pattern. Forefoot striking increases peak stress^16^ and impulse^17,18^ in the Achilles tendon, suggesting an increased injury risk. Forefoot striking also results in higher Achilles tendon forces^19^ and strain rate^16^, potentially indicating greater energy storage, but no studies have estimated the change in energy storage when converting to forefoot striking. In terms of kinematics, forefoot striking is associated with a more plantarflexed ankle^20^ and flexed knee^21^ at initial contact compared to rearfoot striking; these changes result in shorter plantar flexor muscle-tendon lengths and likely shorter muscle fiber lengths, which may affect these muscles’ ability to generate force. Forefoot striking runners produce greater ankle plantarflexion moments during early stance^18,22^, which may indicate greater forces in the plantar flexor muscles. Previous studies have reported increased activity in the gastrocnemius during forefoot striking compared to rearfoot striking^11,23,24^ with no difference in soleus muscle activity^23,25^, suggesting that the gastrocnemius and the soleus may respond differently to forefoot striking. The plantar flexors are crucial during the stance phase of running^9^; thus, it is critical to understand the effect of foot strike pattern on tendon energy storage, the force generation ability of the plantar flexors, and the amount of energy absorbed and generated by these muscles.

The goal of this study was to examine how plantar flexor muscle fiber and tendon mechanics differ between rearfoot and forefoot striking, specifically in the gastrocnemii and soleus. We sought to address four fundamental issues. First, based on the reported effects of forefoot striking on the Achilles tendon^16,19^, we hypothesized that energy storage in the plantar flexor tendons is greater during forefoot striking compared to rearfoot striking. Second, based on differences in knee and ankle kinematics at foot contact between rearfoot and forefoot striking, we anticipated changes to plantar flexor fiber lengths and velocities, and examined how altering foot strike pattern affects the plantar flexor muscles’ ability to generate active force. Third, based on anticipated changes in plantar flexor fiber kinematics and forces, we evaluated how foot strike pattern affects the positive and negative work done by the plantar flexor fibers. Finally, we assessed whether the gastrocnemius and the soleus were affected differently by altering foot strike pattern.

We developed musculoskeletal simulations to characterize the effect of foot strike pattern on plantar flexor muscle-tendon mechanics during running in unprecedented detail (Fig 1; see Methods). Using electromyography (EMG) data and joint kinematics, measured in 16 habitual rearfoot striking subjects running overground using both a rearfoot and forefoot striking pattern, as inputs to our simulations, we simulated the mechanics of the medial gastrocnemius, lateral gastrocnemius and soleus. We analyzed these simulations to evaluate how foot strike pattern affects elastic energy storage in plantar flexor tendons, the force generation ability of the muscles, muscle fiber lengths and velocities, and positive and negative work done by the muscle fibers. These simulations were generated in OpenSim^26^, an open-source simulation software. Experimental data and simulation results are freely available at simtk.org/projects/rfs-ffs-pfs to allow others to reproduce and build upon our work.

**Figure 1.**
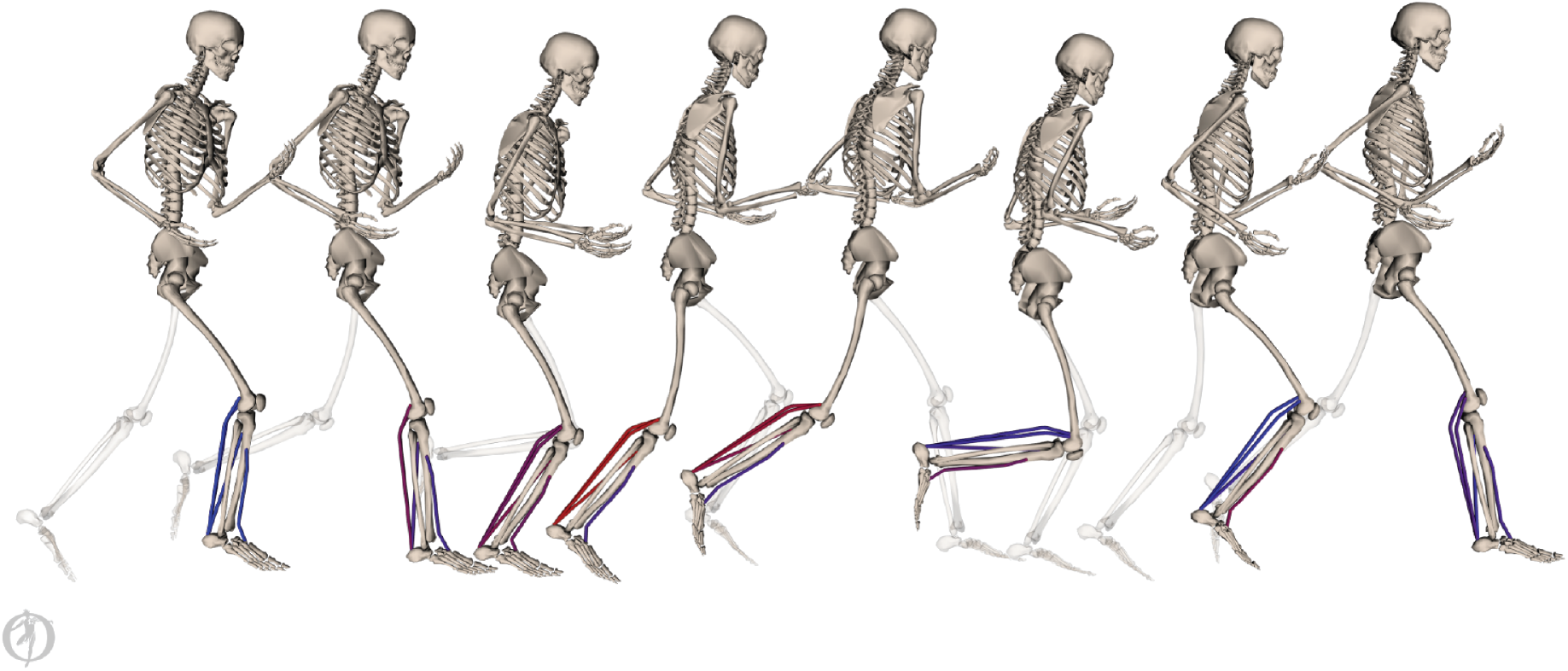
Simulations (pictured) of plantar flexor muscle-tendon mechanics were driven by electromyography data and joint angles. Processed electromyography signals were applied as muscle excitations. Excitations are visualized as a color gradient on the muscles from blue (low excitation) to red (high excitation). Joint angles, estimated from motion capture data, were used to prescribe lower body kinematics in a scaled musculoskeletal model.

## RESULTS

Energy storage in the Achilles tendon was similar (p = 0.703) for rearfoot striking (26.4 ± 4.4 J) and forefoot striking (25.7 ± 7.4 J; Fig 2A). However, foot strike pattern affected tendon energy storage differently for the components of the Achilles tendon associated with the gastrocnemii and the soleus. Energy storage in the gastrocnemius component of the Achilles tendon increased, while energy storage in the soleus component decreased during forefoot striking compared to rearfoot striking (p = 0.002). The timing of peak negative tendon energy storage in the gastrocnemius tendon shifted significantly earlier in the gait cycle during forefoot striking (medial and lateral: p < 0.001), with peak negative tendon power on average shifting from 20% of the gait cycle during rearfoot striking to 6% of the gait cycle during forefoot striking (Fig 2B). The timing for peak negative tendon energy storage in the soleus was not significantly different between rearfoot and forefoot striking (p = 0.487). We estimated tendon energy storage using positive work done by the plantar flexor tendons and assuming no energy loss. Since tendons are modeled as elastic structures, the positive and negative work done by a tendon during the gait cycle should be equivalent. We calculated the difference between positive and negative work done by the tendons and found the average error to be within 0.3 J (2.3%) for all tendons under all conditions. Therefore, we did not test for differences in negative work done by the plantar flexor tendons between foot strike patterns, but expect all differences in positive work to hold for negative work as well.

**Figure 2.**
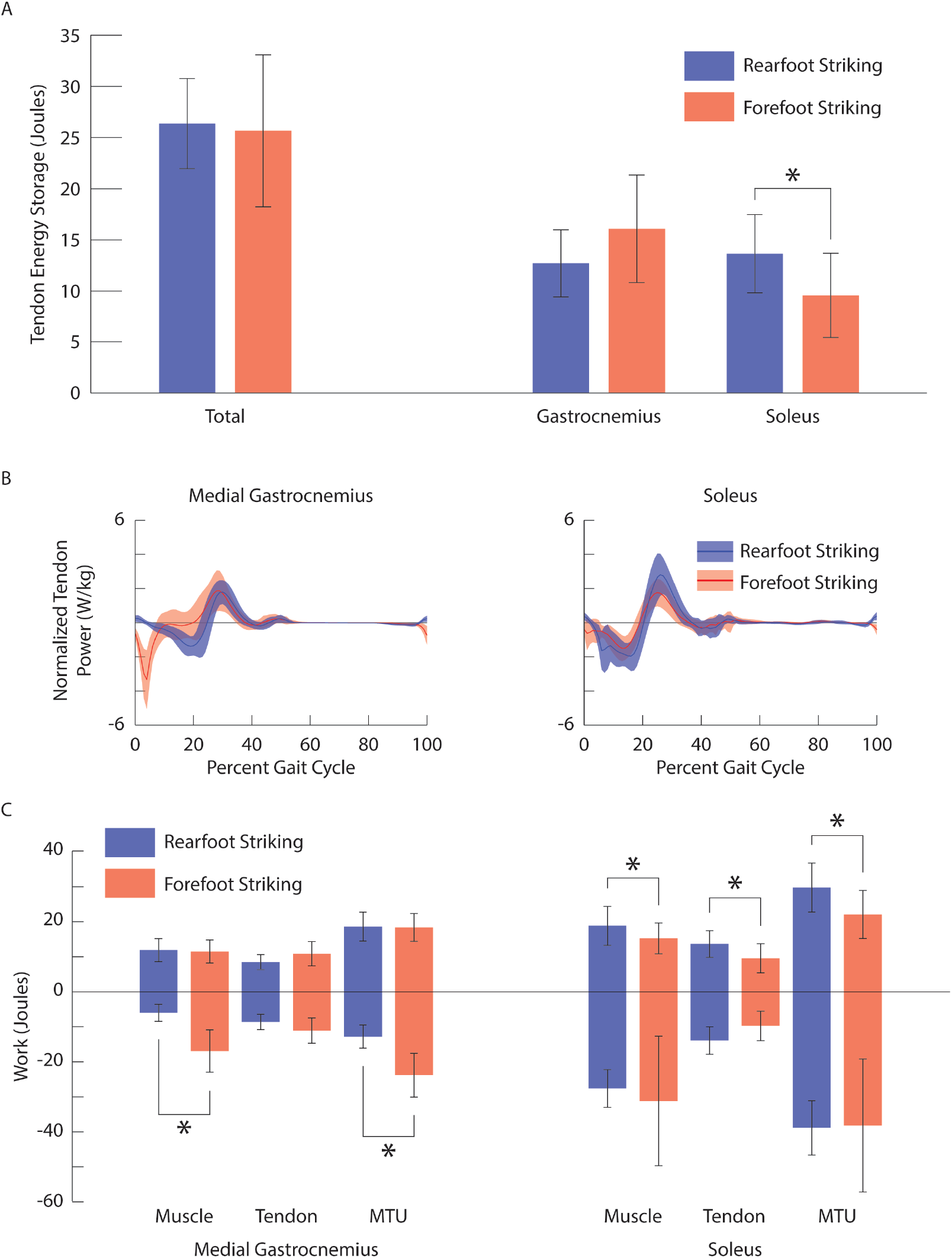
(A) Ensemble average ± one standard deviation tendon energy storage for all plantar flexor tendons together (left), as well as gastrocnemius and soleus components (right) during rearfoot striking (blue) and forefoot striking (red). (B) Ensemble average ± one standard deviation normalized tendon power for the medial gastrocnemius (left) and the soleus (right) during rearfoot striking (blue) and forefoot striking (red). (C) Ensemble average ± one standard deviation positive and negative work done by the muscle fibers, tendon, and muscle-tendon unit of the medial gastrocnemius (left) and the soleus (right) during rearfoot striking (blue) and forefoot striking (red). * indicates a significant difference (p < 0.05).

The force generation ability (i.e. the force generated per unit of activation) at peak active force was higher in the gastrocnemii (medial and lateral: p < 0.001) during forefoot striking compared to rearfoot striking, but was not significantly different in the soleus (p = 0.700; Fig 3; Table 1). Peak active force shifted earlier in the gait cycle for the gastrocnemii during forefoot striking (medial and lateral: p < 0.001), but the timing of peak active force did not significantly change for the soleus (p = 0.061).

**Table 1.** Force generation ability and timing of peak active force for the medial gastrocnemius, lateral gastrocnemius, and soleus during rearfoot striking (RFS) and forefoot striking (FFS). Presented are the mean (standard deviation) and associated p-values. **Bold** indicates a significant difference.

|  | Force Generation Ability |  |  | Timing of Peak Active Force (% Gait Cycle) |  |  |
| --- | --- | --- | --- | --- | --- | --- |
|  | RFS | FFS | p-value | RFS | FFS | p-value |
| Medial Gastrocnemius | 0.79 (0.15) | 1.10 (0.16) | **< 0.001** | 25.7 (2.3) | 16.6 (6.7) | **< 0.001** |
| Lateral Gastrocnemius | 0.80 (0.14) | 1.09 (0.16) | **< 0.001** | 25.9 (2.4) | 16.4 (7.0) | **< 0.001** |
| Soleus | 0.67 (0.12) | 0.69 (0.18) | 0.700 | 21.9 (2) | 19.7 (3.7) | 0.061 |

**Figure 3.**
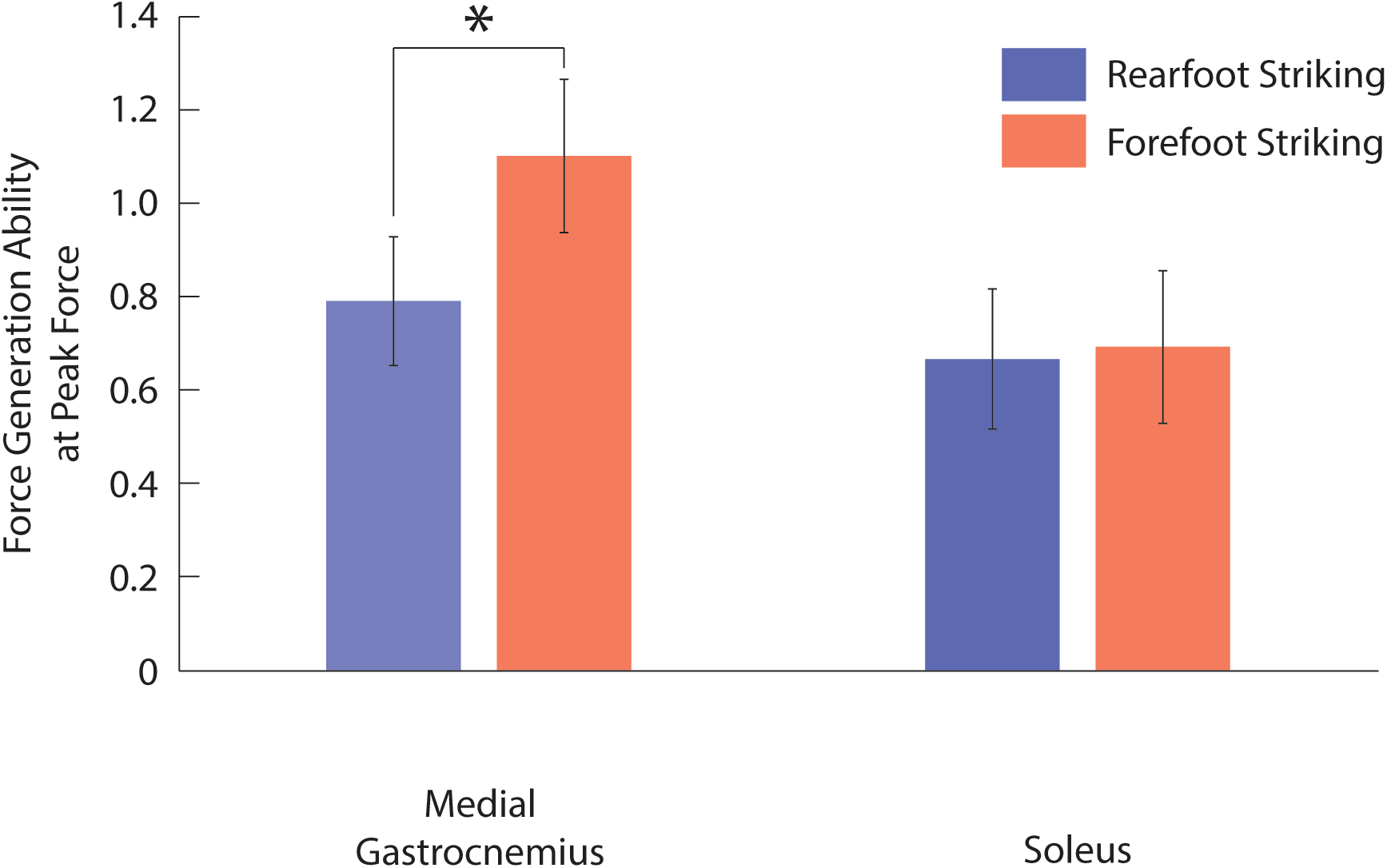
Ensemble average ± one standard deviation force generation ability at peak active force for the medial gastrocnemius (left) and the soleus (right) during rearfoot striking (blue) and forefoot striking (red). * indicates a significant difference (p < 0.05).

Forefoot striking increased activation of the gastrocnemii during 91 – 17% of the gait cycle compared to rearfoot striking (p < 0.001; the same activation was used for both the medial and lateral gastrocnemii; Fig 4), but decreased activation of the soleus during 25 – 34% of the gait cycle (p = 0.014) and 81 – 89% of the gait cycle (p < 0.001). During forefoot striking, gastrocnemii fibers were shorter for the majority of the gait cycle (medial: 80 – 40%, p < 0.001; lateral: 79 – 40%, p < 0.001), and soleus fibers were shorter during 77 – 5% of the gait cycle (p < 0.001) and 30 – 38% of the gait cycle (p = 0.002). Gastrocnemii and soleus fiber lengthening velocities were greater during forefoot striking compared to rearfoot striking early in the stance phase of gait (medial and lateral gastrocnemii: 1 – 9%, p < 0.001; soleus: 0 – 7%, p < 0.001). During 3 – 7% of the gait cycle, the gastrocnemii fibers were lengthening during forefoot striking, but shortening during rearfoot striking. Similarly, during 1 – 4% of the gait cycle, soleus fibers were lengthening during forefoot striking and shortening during rearfoot striking.

**Figure 4.**
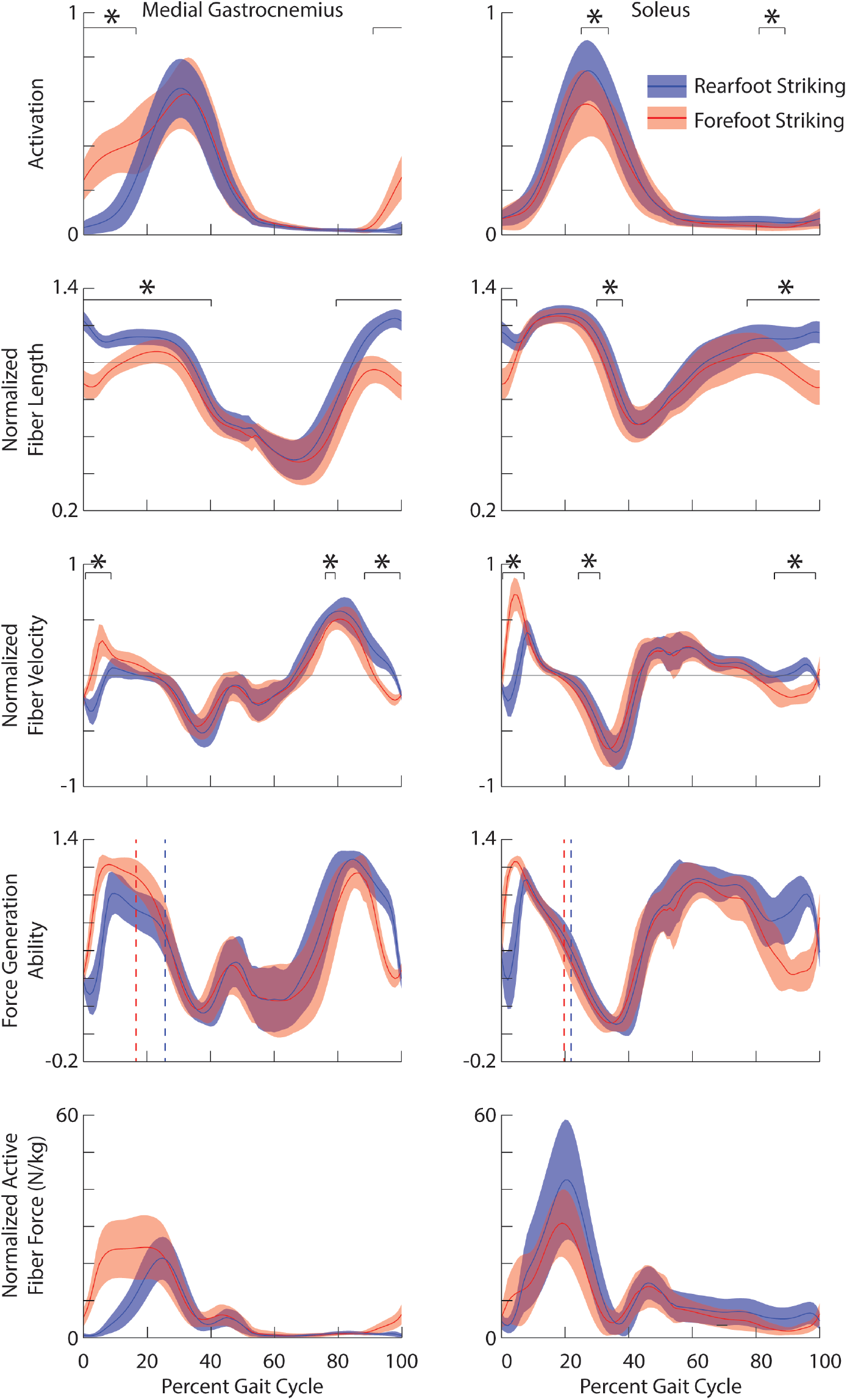
From top to bottom: Ensemble average ± one standard deviation activation, normalized fiber lengths, normalized fiber velocities, force generation ability and normalized active fiber force for medial gastrocnemius (left) and the soleus (right) during rearfoot striking (blue) and forefoot striking (red). Vertical dashed lines represent timing of peak active fiber force. * indicates the portions of the gait cycle when there is a significant difference (p < 0.05).

Converting to forefoot striking resulted in greater negative work done by the gastrocnemii muscle fibers (medial and lateral: p < 0.001) and muscle-tendon units (medial and lateral: p < 0.001), but did not affect the positive work done by the gastrocnemii muscle fibers (medial: p = 0.652; lateral: p = 0.853) or muscle-tendon units (medial: p = 0.723; lateral: p = 0.584; Fig 2C). Conversely, converting to forefoot striking did not significantly affect the negative work done by the soleus muscle fibers (p = 0.480) or muscle-tendon units (p = 0.902), but caused a reduction in the positive work done by the soleus muscle fibers (p = 0.007) and muscle-tendon units (p < 0.001). Our estimates for positive and negative work done by the plantar flexor muscle fibers and muscle-tendon units are presented in Table 2.

**Table 2.** Positive and negative work done by the muscle fibers, tendons, and muscle-tendon units for the lateral gastrocnemius, medial gastrocnemius, and soleus during rearfoot striking (RFS) and forefoot striking (FFS). Presented are the mean (standard deviation) and the associated p-values. **Bold** indicates a significant difference.

|  | Muscle Fiber |  |  | Tendon |  |  | Muscle-Tendon Unit |  |  |
| --- | --- | --- | --- | --- | --- | --- | --- | --- | --- |
|  | RFS | FFS | p-value | RFS | FFS | p-value | RFS | FFS | p-value |
| **Medial Gastrocnemius** |  |  |  |  |  |  |  |  |  |
| Positive Work | 11.9 (3.2) | 11.5 (3.3) | 0.652 | 8.5 (2.2) | 10.8 (3.5) | -- | 18.6 (4.2) | 18.4 (4.0) | 0.723 |
| Negative Work | 6.0 (2.4) | 16.9 (6.0) | **< 0.001** | -- | -- | -- | 12.8 (3.3) | 23.8 (6.3) | **< 0.001** |
| **Lateral Gastrocnemius** |  |  |  |  |  |  |  |  |  |
| Positive Work | 7.1 (1.8) | 7.0 (1.9) | 0.853 | 4.2 (1.1) | 5.2 (1.7) | -- | 10.4 (2.3) | 10.1 (2.1) | 0.584 |
| Negative Work | 3.2 (1.3) | 8.9 (3.2) | **< 0.001** | -- | -- | -- | 6.6 (1.7) | 12.1 (3.3) | **< 0.001** |
| **Soleus** |  |  |  |  |  |  |  |  |  |
| Positive Work | 18.8 (5.6) | 15.3 (4.4) | **0.007** | 13.7 (3.8) | 9.6 (4.1) | **0.002** | 29.8 (7.0) | 22.1 (6.9) | **< 0.001** |
| Negative Work | 27.6 (5.4) | 31.2 (18.5) | 0.480 | -- | -- | -- | 38.8 (7.8) | 38.2 (19.0) | 0.902 |

## DISCUSSION

The purpose of this study was to identify how plantar flexor muscle-tendon mechanics differed between rearfoot and forefoot striking in habitual rearfoot striking runners. We hypothesized that energy storage in the plantar flexor tendons would be greater during forefoot striking yet observed no significant differences in total energy storage between rearfoot and forefoot striking. This occurred because the increase in elastic energy storage in the gastrocnemius tendon was offset by the decrease in elastic energy storage in the soleus tendon. As expected, altering foot strike pattern affected plantar flexor muscle fiber lengths and velocities around foot contact. The changes in plantar flexor fiber kinematics during forefoot striking resulted in increases in the force generation ability of the gastrocnemii at the time it generates peak active force, with no significant effect on the force generation ability of the soleus. When evaluating the work done by the plantar flexor fibers, we found that forefoot striking increased gastrocnemius negative fiber work and decreased positive soleus fiber work. Overall, foot strike pattern affected the gastrocnemii and the soleus muscle-tendon mechanics differently.

Differences in tendon energy storage between the gastrocnemius and the soleus were due, in part, to how forefoot striking affected these muscles’ activation patterns. The activation differences observed during forefoot striking affected the timing for tendon energy storage in the gastrocnemii. Greater activation in the gastrocnemii prior to and immediately after foot contact resulted in greater muscle and tendon forces and, consequently, greater tendon lengthening velocities during forefoot striking. This combination of greater forces and greater lengthening velocities during early stance with forefoot striking caused peak negative tendon power to shift earlier in the gait cycle for the gastrocnemii (Fig 2B). Subjects did not increase activation of the soleus in the early stance phase and experienced a smaller shift in the timing of peak negative tendon power during forefoot striking.

Reviewing changes to force generation ability may provide insight into how increased activation in the gastrocnemii may be beneficial during forefoot striking. Force generation ability represents the muscle’s ability to generate active force, and takes into account the effects of fiber length, fiber velocity and pennation angle^13^. Our simulations show the force generation ability of the soleus at peak active force was not significantly affected by foot strike pattern, while the gastrocnemii had significantly higher force generation ability, as well as higher activation, at the time of peak active force during forefoot striking compared to rearfoot striking (Fig 4). Thus, we postulate that to generate the higher plantar flexion moments found in forefoot striking^22^, runners take advantage of the higher force generation ability of the gastrocnemii during forefoot striking by increasing activation and force in these muscles rather than utilizing the soleus, which does not benefit from improved force generation ability.

Converting to forefoot striking caused runners to increase demands on the gastrocnemii without increasing demand on the soleus. In addition to greater peak muscle forces during forefoot striking, activation was higher in the gastrocnemii after foot contact when the fibers were lengthening. The gastrocnemii fibers were, therefore, undergoing eccentric contraction during forefoot striking compared to concentric contraction during rearfoot striking. Although the soleus fibers were also lengthening after foot contact in forefoot striking compared to shortening in rearfoot striking, we did not find increases in activation, muscle forces or negative work, but instead found a decrease in positive soleus fiber work. The differences we found in muscle-tendon mechanics between the gastrocnemii and the soleus were likely due to knee kinematics and differences in activation between the plantar flexors.

Greater activation, force, and lengthening velocity of the gastrocnemii but not the soleus in forefoot striking has implications for muscle injury and fatigue. Eccentric exercise has previously been shown to improve strength^27^ and potentially, in the long term, prevent injury^28,29^. However, in the short term, eccentric exercise has also been shown to cause muscle soreness, longer-lasting fatigue, and increased risk of muscle damage compared to concentric exercise^30^. While forefoot striking may be beneficial for strengthening the plantar flexors due to increased eccentric contraction, the gastrocnemius may be at increased injury risk in the short-term. Additionally, given that the gastrocnemius is more fatigable than the soleus^31^, higher activation and force in the gastrocnemius may contribute to why runners transition from forefoot striking to rearfoot striking over the course of a long distance run.

While we analyzed habitual rearfoot strikers running with both rearfoot and forefoot striking running patterns, our results may also be applicable to habitual forefoot strikers. Previous work^10^ has shown that acutely trained and habitual forefoot striking runners have similar kinematics. The runners in this study demonstrated increased ankle plantarflexion and knee flexion at initial contact during forefoot striking compared to rearfoot striking^32^, as has been found in habitual forefoot striking runners^20^. Further, the trends in muscle activity are consistent with previously reported differences in muscle activity between habitual rearfoot striking and habitual forefoot striking runners^23^. The similarities in kinematics and muscle activity suggest that our results may also apply to habitual forefoot striking runners.

We modeled the Achilles tendon as three distinct tendons – one for the soleus, one for the medial gastrocnemius and one for the lateral gastrocnemius – as opposed to a single shared tendon. In support of modeling the gastrocnemii and soleus tendons independently, Franz et al.^33^ found that the superficial and deep regions of the Achilles tendon undergo different deformations. Despite modeling distinct tendons, our estimates of total energy storage in the Achilles tendon (rearfoot striking: 26.4 J; forefoot striking: 25.7 J), found by summing the tendon energy storage in all three tendons, were similar to previous estimates of energy storage in the Achilles tendon. Experimental estimates^6,8,34^ ranged from 17-35 J, inverse dynamics estimates^35^ ranged from 10-39 J, and muscle-level simulation estimates^36,37^ ranged from 27-40 J. Our simulations estimated medial gastrocnemius tendon contributions to positive muscle-tendon work of 46%, compared to ultrasound estimates^38^ of 63-70% for rearfoot striking running at similar speeds. This discrepancy may result from modeling choices, particularly tendon compliance. We tested the effect of increasing tendon compliance (from 4.9% to 10% tendon strain at peak force) and found increased energy storage in the Achilles tendon (rearfoot striking: 28.3 J; forefoot striking: 28.4 J) and increased tendon contributions to positive work done by the muscle-tendon units under all conditions. Trends in the effects of foot strike pattern on tendon energy storage were unaffected by a more compliant tendon in our simulations.

Although our trends in fiber kinematics are mostly consistent with the literature, our reported fiber lengths are longer than previous estimates. During both rearfoot and forefoot striking, estimated soleus fibers exhibited a smaller fascicle excursion than the gastrocnemii fibers, consistent with previous ultrasound work ^12,14^. The shorten-stretch-shorten kinematics of our estimated gastrocnemius fibers during forefoot striking (Fig 4) are consistent with Ishikawa and Komi^39^ who studied forefoot striking running. However, our gastrocnemius fiber velocities estimated during rearfoot striking, which were mostly negative with relatively small positive velocities, do not agree with reported medial gastrocnemius fibers shortening throughout stance^15^. Our results may differ due to our subjects running at faster speeds and running overground rather than on a treadmill. Previous simulations^13,36^ and ultrasound imaging^39^ estimated fiber lengths to be less than one optimal fiber length throughout the gait cycle when running at similar speeds, but in our simulations peak fiber length exceeded one optimal fiber length. Differences in normalized fiber lengths likely result from using a different model, different plantar flexor tendon compliance, and different estimates of optimal fiber lengths. Recent studies have shown that sarcomeres in healthy individuals are longer than previously estimated using simulation^40,41^, which supports our estimates of longer normalized fiber lengths. Although we may overestimate normalized fiber lengths of the plantar flexors, the paired nature of our data lends confidence to our results demonstrating the effect of foot strike pattern on normalized fiber lengths in running.

This study identified differences in plantar flexor tendon energy storage, and the positive and negative work done by the plantar flexor muscle fibers and muscle-tendon units between rearfoot and forefoot striking. Although positive work done by the soleus fibers decreased, negative work done by the gastrocnemius fibers was higher in forefoot striking compared to rearfoot striking. We reported differences in plantar flexor fiber lengths and fiber velocities between foot strike patterns, and showed that during peak active force, the gastrocnemii fibers were in a more favorable state for generating forces. We postulate that converted forefoot striking runners make use of the improved state of the gastrocnemius by activating this muscle rather than the soleus, a trend that we believe extends to habitual forefoot strikers. Overall, this research supports the notion that runners considering transitioning from rearfoot striking to forefoot striking may benefit from a progressive eccentric strengthening program targeting the plantar flexors to prepare them for the increased demands of forefoot striking and to help avoid injury.

## METHODS

### Experimental Data

We collected data from 16 habitual rearfoot striking subjects running overground using both a rearfoot and forefoot striking pattern. The subjects were healthy recreational runners who reported running at least 10 km per week (11 females, 5 males; age: 32.1 ± 9.9 years; height: 167 ± 10 cm; mass: 62.5 ± 10.1 kg; mileage: 36.1 ± 19.5 km/week). The subjects ran at their self-selected speed (2.94 ± 0.30 m/s) using their habitual rearfoot striking pattern and a forefoot striking pattern after acute gait retraining with visual feedback as described previously^32^.

We tracked the positions of 43 reflective markers from a full-body marker set at 200 Hz using a motion capture system (Motion Analysis Corporation, Santa Rosa, CA, USA) and collected ground reaction forces at 2000 Hz using in-ground force plates (Bertec Corporation, Columbus, OH, USA). Simultaneously, we collected surface EMG data (Delsys Inc., Boston, MA, USA) from the medial gastrocnemius, soleus and tibialis anterior at 2000 Hz. For both rearfoot and forefoot striking, we analyzed data from three trials, each of which captured one stride from each subject’s dominant limb. Each subject gave informed consent prior to participation according to a protocol approved by the Stanford University Institutional Review Board.

### Musculoskeletal Model

We used a full-body musculoskeletal model with 29 degrees of freedom^42^. The model included six degrees of freedom to position and orient the pelvis in space, three-degree-of-freedom ball-and-socket joints to represent each hip joint, custom one-degree-of-freedom joints to represent each knee joint and one-degree-of-freedom revolute joints to represent each ankle joint. The model included 40 muscles per lower limb, but this study focused on analyzing only the soleus, medial gastrocnemius and lateral gastrocnemius of the dominant limb. Muscle-tendon units were modeled using Hill-type muscle models as described by Millard et al.^43^.

### Simulation of Muscle-tendon Dynamics

We started by scaling the generic musculoskeletal model to each subject’s anthropometry. In addition to scaling body segments, we scaled muscle-tendon parameters such that the ratio of optimal fiber length and tendon slack length was preserved. We then used OpenSim’s inverse kinematics algorithm^26^ to calculate joint angles that best tracked subjects’ measured marker positions during the running trials.

Muscle excitations were applied to the plantar flexors and the tibialis anterior based on surface EMG data. Data from the medial gastrocnemius, soleus, and tibialis anterior were filtered using a band-pass filter (50 – 500Hz), rectified and filtered again using a critically damped low-pass filter with a 15 Hz cutoff. The filtered EMG signals were scaled to peak activity in the muscle over all running trials. We then applied a 40 ms delay to account for electromechanical delay, consistent with previous work^13^. Processed EMG signals were applied as excitations to the plantar flexors and the tibialis anterior, with the medial gastrocnemius signal applied to both the medial and lateral gastrocnemii in the model. Previous studies have shown measured muscle activity to be similar between the medial and lateral gastrocnemii during both rearfoot and forefoot striking^13,23,44^, and other simulation studies have used medial gastrocnemius activity to define excitations for both the medial and lateral gastrocnemii^37^. For all other muscles, we applied an excitation of 0.01 throughout the simulation.

Joint angles, estimated from inverse kinematics, and muscle excitations, derived from processed EMG data, drove forward simulations of muscle-tendon dynamics (Fig 5). From these simulations, we estimated normalized fiber lengths, normalized fiber velocities, and fiber forces, along with power done by the muscle-tendon units, muscle fibers, and tendons. Using these results, we estimated the force generation ability^13^ of each plantar flexor muscle throughout the gait cycle. We also estimated positive and negative work done by the plantar flexor muscle-tendon units and muscle fibers. Tendon energy storage was estimated using positive work done by the plantar flexor tendons and assuming no energy loss. We estimated Achilles tendon energy storage by summing the positive work done by the gastrocnemii and soleus tendons. We estimated the gastrocnemius component of Achilles tendon energy storage by summing the positive work done by the medial and lateral gastrocnemii tendons. To estimate positive and negative work, we integrated the positive and negative parts of the power curves, respectively. The library of these simulations is publicly available at simtk.org/projects/rfs-ffs-pfs.

**Figure 5.**
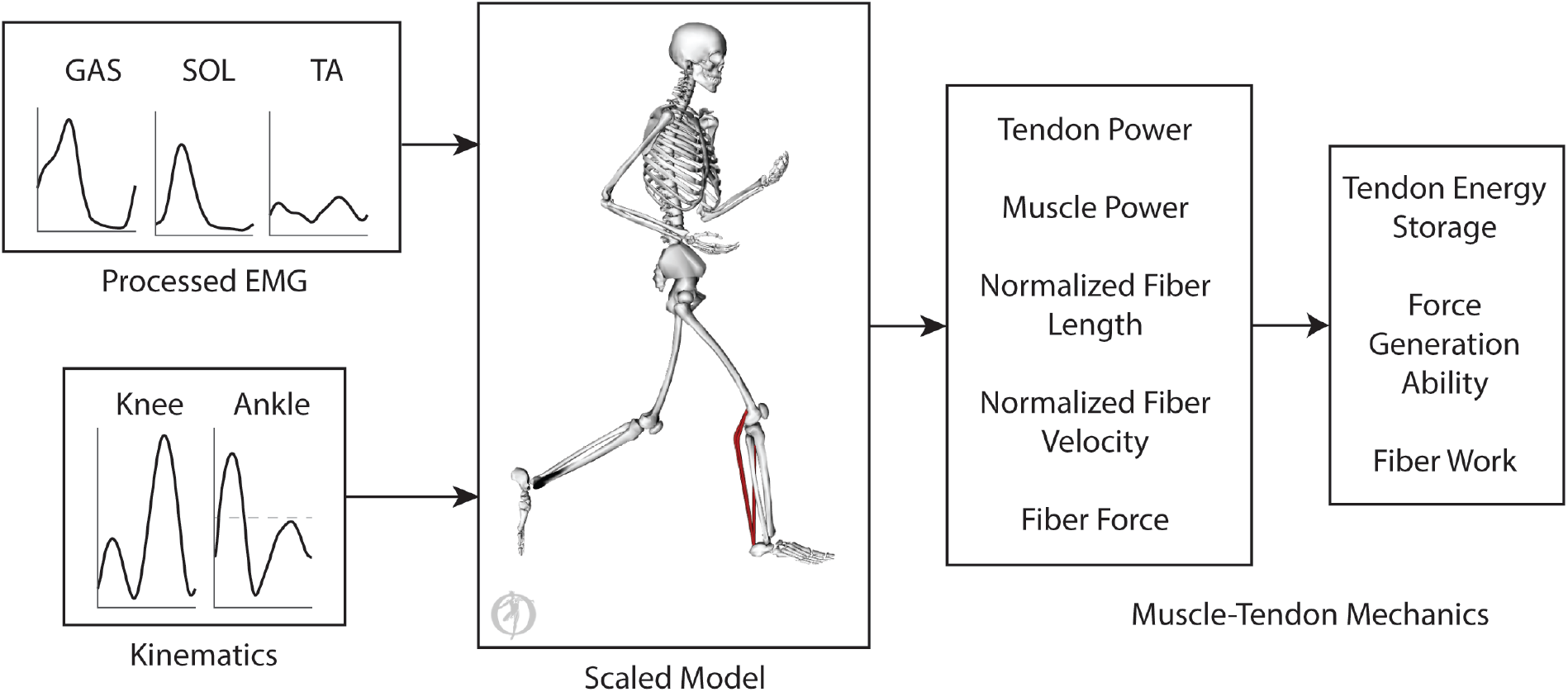
Simulations of plantar flexor muscle-tendon mechanics were driven by electromyography data, applied as muscle excitations, and joint angles, which prescribed lower body kinematics. The simulations were used to estimate plantar flexor tendon power, muscle power, normalized fiber lengths, normalized fiber velocities and fiber forces. Plantar flexor tendon energy storage and positive and negative fiber work were estimated by integrating tendon power and muscle power. Force generation ability, which combines the effects of fiber length, fiber velocity and pennation angle, was also estimated.

### Testing the simulations by comparison to experimental data

To validate our simulations, we compared the sum of ankle moments and powers produced by the plantar flexors and the tibialis anterior from our forward simulations with the net ankle moment and power estimated from inverse dynamics during stance (Fig 6). Although our solutions only included excitations from three plantar flexor muscles and the tibialis anterior, the soleus along with the medial and lateral gastrocnemii can produce 93% of the model’s maximum plantarflexion moment, and the tibialis anterior can produce 62% of the model’s maximum dorsiflexion moment.

**Figure 6.**
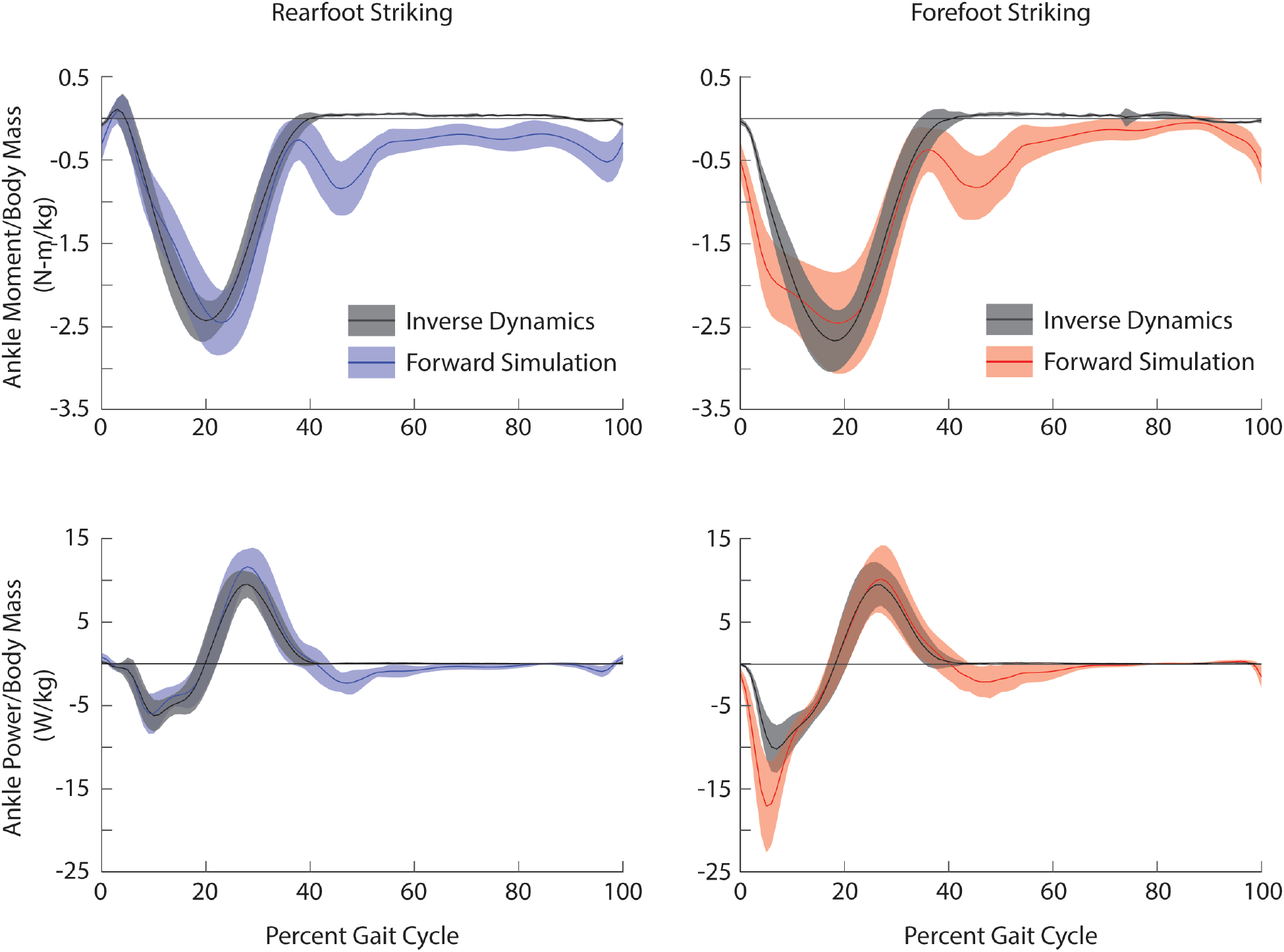
Ensemble average ± one standard deviation simulated ankle joint moments (top) and powers (bottom) estimated from forward simulations and inverse dynamics during rearfoot striking (left) and forefoot striking (right) during running. During the forward simulations, average simulated ankle joint moments were estimated based on contributions from the tibialis anterior, the medial and lateral gastrocnemii, and the soleus. Average ankle power was estimated by multiplying the summed moment with the ankle angular velocity. Forward simulation results for rearfoot and forefoot striking are shown in blue and red, respectively. Inverse dynamics results are shown in black.

When comparing peak plantarflexion moments between the forward simulation and inverse dynamics results, the average timings were within 3% of the gait cycle and the average magnitudes were within one standard deviation during both rearfoot and forefoot striking. When comparing the average timings for peak positive and peak negative ankle power, the forward simulation and inverse dynamics results were within 2% of the gait cycle during both rearfoot and forefoot striking. For the rearfoot striking forward simulations, the magnitude of peak negative ankle power was within one standard deviation and the magnitude of peak positive ankle power was within two standard deviations of the inverse dynamics results. For the forefoot striking forward simulations, the magnitude of peak negative ankle power was within three standard deviations and the magnitude of peak positive ankle power was within one standard deviation of the inverse dynamics results. Aside from peak negative ankle power during forefoot striking, our errors were within the guidelines of two standard deviations, as recommended by Hicks et al.^45^. We tested the importance of our mismatch in peak negative ankle power during forefoot striking by adjusting our model to better match these peaks. We were able to best align the peak negative ankle powers during forefoot striking by scaling our muscle activity such that peak excitation was 0.8 and, as has been done in previous studies^13,46^, increasing tendon compliance in the plantar flexors to 10% strain at maximum isometric force. We ultimately chose not to use these adjustments due to obvious mismatches in peak ankle moments during both rearfoot and forefoot striking, and peak negative ankle power during rearfoot striking.

### Statistical Analysis

We compared plantar flexor tendon energy storage, the timing of plantar flexor tendon energy storage, the positive and negative work done by the plantar flexor muscle-tendon units and muscle fibers, the force generation ability of the plantar flexors at peak active force, and the timing of peak active force using paired t-tests. These analyses were done using SPSS (SPSS, IBM, Armonk, NY, USA) and significance for all analyses, before corrections, was set at p < 0.05.

To identify portions of the gait cycle when fiber lengths and velocities were significantly different between foot strike patterns, we compared the trajectories of plantar flexor normalized fiber lengths and normalized fiber velocities using statistical parametric mapping^47^. This method was also used to compare how plantar flexor activations differ between foot strike patterns. Statistical parametric mapping was designed to identify time ranges when continuous curves are significantly different. While testing for differences in our curves, we indicated that the data were paired, included wrapping since our data are cyclical, included Bonferroni corrections when necessary, and set significance, before corrections, at p < 0.05. These analyses were done using “SPM1D” (version M.0.4.5, www.spm1d.org), a free and open source software package for statistical parametric mapping in Matlab (R2015b, The Mathworks Inc., Natick, MA, USA). We excluded any time ranges less than 1% of the gait cycle that were identified as significantly different because our data does not have sufficient resolution to detect such changes.

## CONFLICT OF INTEREST STATEMENT

None of the authors had any financial or personal conflict of interest with regard to this study.

## ACKNOWLEDGEMENTS

The authors thank Carmichael Ong, Apoorva Rajagopal, and Johann Simpson. The authors also thank Irene Davis for the early conversations regarding this project. This work was supported by a Stanford Bio-X Graduate Student Fellowship, a grant from the PAC 12, and NIH grants U54 EB020405, P2C HD065690.

## AUTHOR CONTRIBUTIONS

All authors contributed to the study formation; J.Y. implemented the analysis and drafted the manuscript and figures; C.D. helped develop tools for the analysis; A.S. contributed to the acquisition of data; all authors reviewed and edited the manuscript and figures.

## Notes

https://simtk.org/projects/rfs-ffs-pfs

## REFERENCES

1. Blickhan, R. The spring-mass model for running and hopping. J. Biomech. 22, 1217–1227 (1989).

2. Bullimore, S. R. & Burn, J. F. Ability of the planar spring-mass model to predict mechanical parameters in running humans. J. Theor. Biol. 248, 686–695 (2007).

3. Dalleau, G., Belli, A., Bourdin, M. & Lacour, J. R. The spring-mass model and the energy cost of treadmill running. Eur. J. Appl. Physiol. Occup. Physiol. 77, 257–263 (1998).

4. Morin, J. B., Jeannin, T., Chevallier, B. & Belli, A. Spring-mass model characteristics during sprint running: Correlation with performance and fatigue-induced changes. Int. J. Sports Med. 27, 158–165 (2006).

5. Girard, O., Micallef, J. P. & Millet, G. P. Changes in spring-mass model characteristics during repeated running sprints. Eur. J. Appl. Physiol. 111, 125–134 (2011).

6. Alexander, R. M. & Bennet-Clark, H. C. Storage of elastic strain energy in muscle and other tissues. Nature 265, 114–117 (1977).

7. Winter, D. A. Moments of force and mechanical power in jogging. J. Biomech. 16, 91–97 (1983).

8. Ker, R. F., Bennett, M. B., Bibby, S. R., Kester, R. C. & Alexander, R. M. The spring in the arch of the human foot. Nature 325, 147–149 (1987).

9. Hamner, S. R., Seth, A. & Delp, S. L. Muscle contributions to propulsion and support during running. J. Biomech. 43, 2709–2716 (2010).

10. Williams, D., McClay, I. & Manal, K. Lower extremity mechanics in runners with a converted forefoot strike pattern. J. Appl. Biomech. 16, 210–218 (2000).

11. Giandolini, M. et al. Impact reduction during running: efficiency of simple acute interventions in recreational runners. Eur. J. Appl. Physiol. 113, 599–609 (2013).

12. Cronin, N. J. & Finni, T. Treadmill versus overground and barefoot versus shod comparisons of triceps surae fascicle behaviour in human walking and running. Gait Posture 38, 528–533 (2013).

13. Arnold, E. M., Hamner, S. R., Seth, A., Millard, M. & Delp, S. L. How muscle fiber lengths and velocities affect muscle force generation as humans walk and run at different speeds. J. Exp. Biol. 216, 2150–2160 (2013).

14. Lai, A. K. M., Lichtwark, G. A., Schache, A. G. & Pandy, M. G. Differences in in vivo muscle fascicle and tendinous tissue behavior between the ankle plantarflexors during running. Scand. J. Med. Sci. Sport. 1828–1836 (2018). doi:10.1111/sms.13089

15. Lichtwark, G. A., Bougoulias, K. & Wilson, A. M. Muscle fascicle and series elastic element length changes along the length of the human gastrocnemius during walking and running. J. Biomech. 40, 157–164 (2007).

16. Lyght, M., Nockerts, M., Kernozek, T. W. & Ragan, R. Effects of foot strike and step frequency on Achilles tendon stress during running. J. Appl. Biomech. 32, 365–372 (2016).

17. Almonroeder, T., Willson, J. D. & Kernozek, T. W. The effect of foot strike pattern on achilles tendon load during running. Ann. Biomed. Eng. 41, 1758–1766 (2013).

18. Rice, H. & Patel, M. Manipulation of Foot Strike and Footwear Increases Achilles Tendon Loading during Running. Am. J. Sports Med. 45, 2411–2417 (2017).

19. Kulmala, J. P., Avela, J., Pasanen, K. & Parkkari, J. Forefoot strikers exhibit lower running-induced knee loading than rearfoot strikers. Med. Sci. Sports Exerc. 45, 2306–2313 (2013).

20. Lieberman, D. E. et al. Foot strike patterns and collision forces in habitually barefoot versus shod runners. Nature 463, 531–535 (2010).

21. Laughton, C., Davis, I. & Hamill, J. Effect of strike pattern and orthotic intervention on tibial shock during running. J. Appl. Biomech. 19, 153–168 (2003).

22. Rooney, B. D. & Derrick, T. R. Joint contact loading in forefoot and rearfoot strike patterns during running. J. Biomech. 46, 2201–2206 (2013).

23. Yong, J. R., Silder, A. & Delp, S. L. Differences in muscle activity between natural forefoot and rearfoot strikers during running. J. Biomech. 47, 3593–3597 (2014).

24. Shih, Y., Lin, K. L. & Shiang, T. Y. Is the foot striking pattern more important than barefoot or shod conditions in running? Gait Posture 38, 490–494 (2013).

25. Landreneau, L. L., Watts, K., Heitzman, J. E. & Childers, W. L. Lower limb muscle activity during forefoot and rearfoot strike running techniques. Int. J. Sports Phys. Ther. 9, 888–97 (2014).

26. Delp, S. L. et al. OpenSim: open-source software to create and analyze dynamic simulations of movement. IEEE Trans. Biomed. Eng. 54, 1940–1950 (2007).

27. Johnson, B. L. Eccentric vs concentric muscle training for strength development. Medicine and Science in Sports 4, 111–115 (1972).

28. Gabbe, B. J., Branson, R. & Bennell, K. L. A pilot randomised controlled trial of eccentric exercise to prevent hamstring injuries in community-level Australian Football. J. Sci. Med. Spor. 9, 103–109 (2006).

29. Petersen, J., Thorborg, K., Nielsen, M. B., Budtz-Jørgensen, E. & Hölmich, P. Preventive effect of eccentric training on acute hamstring injuries in Men’s soccer: A cluster-randomized controlled trial. Am. J. Sports Med. 39, 2296–2303 (2011).

30. Clarkson, P. M. & Newham, D. J. Associations between muscle soreness, damage, and fatigue. Adv Exp Med Biol 384, 457–469 (1995).

31. Ochs, R. M., Smith, J. L. & Edgerton, V. R. Fatigue characteristics of human gastrocnemius and soleus muscles. Electromyogr. Clin. Neurophysiol. 17, 297–306 (1977).

32. Yong, J. R., Silder, A., Montgomery, K. L., Fredericson, M. & Delp, S. L. Acute changes in foot strike pattern and cadence affect running parameters associated with tibial stress fractures. J. Biomech. (2018). doi:10.1016/j.jbiomech.2018.05.017

33. Franz, J. R., Slane, L. C., Rasske, K. & Thelen, D. G. Non-uniform in vivo deformations of the human Achilles tendon during walking. Gait Posture 41, 192–197 (2015).

34. Kyröläinen, H., Finni, T., Avela, J. & Komi, P. V. Neuromuscular behaviour of the triceps surae muscle-tendon complex during running and jumping. Int. J. Sports Med. 24, 153–155 (2003).

35. Hof, A. L., Van Zandwijk, J. P. & Bobbert, M. F. Mechanics of human triceps surae muscle in walking, running and jumping. Acta Physiol. Scand. 174, 17–30 (2002).

36. Lai, A., Schache, A. G., Lin, Y.-C. & Pandy, M. G. Tendon elastic strain energy in the human ankle plantar-flexors and its role with increased running speed. J. Exp. Biol. 217, 3159–68 (2014).

37. Sasaki, K. & Neptune, R. R. Muscle mechanical work and elastic energy utilization during walking and running near the preferred gait transition speed. Gait Posture 23, 383–390 (2006).

38. Farris, D. J. & Sawicki, G. S. Human medial gastrocnemius force–velocity behavior shifts with locomotion speed and gait. Proc. Natl. Acad. Sci. 109, 977–982 (2012).

39. Ishikawa, M. & Komi, P. V. The role of the stretch reflex in the gastrocnemius muscle during human locomotion at various speeds. J. Appl. Physiol. 103, 1030–1036 (2007).

40. Chen, X. & Delp, S. L. Human soleus sarcomere lengths measured using in vivo microendoscopy at two ankle flexion angles. J. Biomech. 49, 4164–4167 (2016).

41. Son, J., Indresano, A., Sheppard, K., Ward, S. R. & Lieber, R. L. Intraoperative and biomechanical studies of human vastus lateralis and vastus medialis sarcomere length operating range. J. Biomech. 67, 91–97 (2018).

42. Rajagopal, A. et al. Full body musculoskeletal model for muscle-driven simulation of human gait. IEEE Trans. Biomed. Eng. 63, 2068–2079 (2016).

43. Millard, M., Uchida, T., Seth, A. & Delp, S. L. Flexing Computational Muscle: Modeling and Simulation of Musculotendon Dynamics. J. Biomech. Eng. 135, 021005 (2013).

44. Hamner, S. R. & Delp, S. L. Muscle contributions to fore-aft and vertical body mass center accelerations over a range of running speeds. J. Biomech. 46, 780–787 (2013).

45. Hicks, J. L., Uchida, T. K., Seth, A., Rajagopal, A. & Delp, S. L. Is My Model Good Enough? Best Practices for Verification and Validation of Musculoskeletal Models and Simulations of Movement. J. Biomech. Eng. 137, 020905 (2015).

46. Jackson, R. W., Dembia, C. L., Delp, S. L. & Collins, S. H. Muscle–tendon mechanics explain unexpected effects of exoskeleton assistance on metabolic rate during walking. J. Exp. Biol. 220, 2082–2095 (2017).

47. Pataky, T. C. Generalized n-dimensional biomechanical field analysis using statistical parametric mapping. J. Biomech. 43, 1976–1982 (2010).

